# Early endocytosis as a key to understanding mechanisms of plasmalemma tension regulation in filamentous fungi

**DOI:** 10.1101/2020.05.18.102947

**Authors:** Igor Mazheika, Oxana Voronko, Olga Kamzolkina

**Affiliations:** Lomonosov Moscow State University, Moscow, Russia, 119991; Vavilov Institute of General Genetics, Russian Academy of Sciences, Moscow, Russia, 117971

**Keywords:** actin cytoskeleton, basidiomycetes, curtain model, cytochalasin D, latrunculin A, *Rhizoctonia solani*, tubular invaginations, turgor pressure

## Abstract

Two main systems regulate the plasmalemma tension and provide a close connection of the protoplast with the cell wall in fungi: turgor pressure and actin cytoskeleton. These systems work together with the plasmalemma focal adhesion to the cell wall and their contribution to fungal cell organization has been partially studied, but remains controversial in model filamentous ascomycetes and oomycetes, and even less investigated in filamentous basidiomycetes. Early endocytosis, in which F-actin is actively involved, can be used to research of mechanisms regulating the plasmalemma tension, since the latter influences on the primary endocytic vesicles formation. This study examined the effects of actin polymerization inhibitors and hyperosmotic shock on early endocytosis and cell morphology in two filamentous basidiomycetes. The main obtained results: (i) depolymerization of F-actin leads to the fast formation of primary endocytic vesicles but to inhibition of their scission; (ii) moderate hyperosmotic shock does not affect the dynamics of early endocytosis. These and a number of other results allowed offering a curtain model of regulation the plasmalemma tension in basidiomycetes. According to this model, the plasmalemma tension in many nonapical cells of hyphae is more often regulated not by turgor pressure, but by a system of actin driver cables that are associated with the proteins of focal adhesion sites. The change in the plasmalemma tension occurs similar to the movement of the curtain along the curtain rod using the curtain drivers. This model addresses the fundamental properties of the fungal structure and physiology and requires confirmation, including through the yet technically unavailable high quality labeling of the actin cytoskeleton of basidiomycetes.

## INTRODUCTION

Endocytosis is the highly important process by which a living cell uptakes liquid, molecules and particles from the environment and components from its own cytoplasmic membrane. At the initial stages of endocytosis, the formation of a primary endocytic vesicle (PEV) occurs from a limited site of the plasmalemma. Various mechanisms of PEV formation and their scission from the cytoplasmic membrane have been suggested (Johannes et al. 2016; Hinze and Boucrot 2018; Renard et al. 2018; Sandvig et al. 2018). PEVs usually have a round or a rounded elongated shape with a diameter from tens of nanometers to a few micrometers; the membrane of large vesicle can be plicated (with secondary invaginations) (Doherty and McMahon 2009; Hinze and Boucrot 2018). Endocytic plasmalemma invagination can also be as a long narrow tube (PET – primary endocytic tube; synonym for PEV and PET – endocytic carrier). In some cases, PET is a stable formation inherent in a certain type of endocytosis (FEME, for example) (Hinze and Boucrot 2018); but more often, PET results from vesicle scission delay from the plasmalemma with active elongation of the neck of the membrane invagination for some causes. In experimental studies, the formation of PETs is artificially stimulated by various methods (by actin depolymerization and hyperosmotic shock, overexpression of BAR proteins, suppression of dynamin, exposition in GTP-γS, etc.) (Takei et al. 1995; Lee et al. 2002; Itoh et al. 2005; Fricke et al. 2009; Chang-Ileto et al. 2011; Mooren et al. 2012).

There are three universal and conserved participants of the early stages of endocytosis, regardless of the species of organism and type of endocytosis itself: specific membrane lipids (mainly phosphatidylinositol), BAR proteins (proteins that have a BAR domain, for example, amphiphysins, endophilins, GRAFs, SNXs, and many others) and F-actin.

Phosphatidylinositol, due to different variants of the arrangement of phosphate groups in its molecule, provides a signal code for triggering one or another stage or type of endocytosis (Posor et al. 2015). BAR proteins are the “brain” of the initial stages of endocytosis; they control recruitment to the invagination site and the subsequent regulation of many proteins: adaptor and coat proteins, dynamin and numerous others (Itoh et al. 2005; Renard et al. 2018). They can trigger the inhibition or polymerization of actin structures associated with plasmalemma invagination. In addition to their control functions, BAR proteins can directly participate in the initial curving of the membrane during the formation of PEVs, in narrowing and elongating the neck of PEVs, and in PEV scission (for example, in FDS – friction-driven scission) (Hinze and Boucrot 2018; Renard et al. 2018).

Microfilaments of F-actin form two types of basic structures in a living cell: actin networks and actin bundles/cables (Berepiki et al. 2011; Hinze and Boucrot 2018). Actin networks are formed due to Arp2/3-dependent branching of microfilaments. The networks can be cross-linked by specific proteins and form strong and relatively rigid structures. An example is the actin cortex in animal cells that lining the inner surface of the cell and having an average thickness about 200 nm (Clark et al. 2013). Another example is fungal actin patches – scaffolds around PEVs formed by local cross-linked Arp2/3 actin networks. The presence of the actin cortex in fungi is not shown. In addition to the patches, actin rings and cables were found in fungi. These rings and cables are formed formin-dependent due to the parallel connection and cross-linking of actin microfilaments (Berepiki et al. 2011; Kilaru et al. 2015).

Involvement of F-actin in the initial stages of endocytosis in living organisms can be different (see SUPPLEMENTARY FIG. 1A–L). Actin most often performs a force function in early endocytosis, together with myosin and other actin-binding proteins, forming intracellular structures that create traction force and mechanical support for the formation and scission of PEVs (in some fungi, the main such structure is actin patch) (Berepiki et al. 2011; Ferreira and Boucrot 2018). The force that must be applied to invaginate the plasmalemma, and which actin provides, depends as minimum on the endocytic cargo weight and the tension of the plasmalemma. The plasmalemma tension (PMT) can be regulated in different ways. In animal cells, for example, this tension may depend on the rigidity of actin cortex and/or on certain external conditions, on the state and type of cells, the stage of the cell cycle, etc. (for example, when a cell adheres to and spreads on a substrate surface, the plasmalemma tension can increase) (Boulant et al. 2011; Mooren et al. 2012; Kozlov and Chernomordik 2015).

Early endocytosis is indicative to study mechanisms of PMT regulation, especially if using actin polymerization inhibitors (APIs) such as latrunculins and cytochalasins. For example, in fission yeast with relatively high PMT due to turgor pressure, even a small violation of F-actin can lead to a complete blockage of endocytosis (PEVs are not formed; Basu et al. 2014). With weak PMT BAR proteins with other proteins are able to form PEVs (and often PETs) without significant involvement of actin (Mooren et al. 2012; Hinze and Boucrot 2018). Therefore, in budding yeast with full depolymerized F-actin, hyperosmotic condition decreasing PMT can partially restore early endocytosis (Aghamohammadzadeh, Ayscough 2009).

Two main factors affect PMT in organisms with a cell wall, in particular in plants, fungi and fungi-like organisms: turgor pressure and actin cytoskeleton. In addition, there is a third factor in the organisms with a cell wall, which in itself probably has little effect on PMT, but retains the cell integrity at low PMT values. This third factor is focal adhesion of the plasmalemma to the cell wall. The plasmalemma is adhered to the cell wall at plural focal adhesion cites (FACs) using transmembrane integrin-like or other proteins (Oparka 1994; Heath, Steinberg 1999; Chitcholtan, Garrill 2005; Elhasi, Blombers 2019). The intracellular domains of these proteins can be associated with F-actin, and the outer domains are bond with glucans and proteins of the cell wall. FACs also serve to transmit signals from the cell wall and plasmalemma to the cytoplasma compartments (Chitcholtan, Garrill 2005; Elhasi, Blombers 2019). For example, in fungi, removal of the cell wall or osmotic shock serve as a signal for disassembling actin cables inside the cells (SUPPLEMENTARY TABLE 1). It is generally accepted that in fungi, the connection between the plasmalemma and the cell wall is especially close (Walker, White 2017; about the peculiar properties of plasmolysis in yeast and filamentous fungi, see SUPPLEMENTARY TABLE 1).

From SUPPLEMENTARY TABLE 1 it is seen that in the studied groups of fungi and fungi-like organisms, the contribution of turgor and actin cytoskeleton in generation and regulation of PMT is different. In yeast, the main factor affecting PMT is turgor pressure (Basu et al. 2014) and, apparently, it should not be necessarily high. In filamentous ascomycetes, turgor is most likely important for apical and some other cells (Lew et al. 2004; Abadeh, Lew 2013; Lew 2019). In other parts of the mycelium, the cytoskeleton may compete with turgor for influence on PMT. In oomycetes, F-actin as a factor determining PMT can dominate even in apical cells (Walker et al. 2006).

In filamentous basidiomycetes, issues related to the regulation of PMT are less studied than in ascomycetes. In our previous papers (Kamzolkina et al. 2017; Mazheika et al. 2020) two features of early endocytosis in filamentous basidiomycetes was described which may shed light on the mechanism of regulation of PMT in these fungi. First, it has been shown that filamentous fungi have at least two types of endocytosis, differing in the maximum size of PEVs. In addition to the patch type of endocytosis with small-sized PEVs, which is characteristic of many model ascomycetes also macrovesicular type of endocytosis exists. Macrovesicular endocytosis was found in a number of xylotrophic basidiomycetes and in *Coprinus comatus*. The size of PEVs with this type of endocytosis reaches 2 μm (Mazheika et al. 2020). Obviously, such large invaginations of the plasmalemma are not compatible with high turgor pressure. Second, it was preliminarily observed that treatment of the mycelium of *Rhizoctonia solani* and *Stereum hirsutum* by APIs leads to the result described in SUPPLEMENTARY TABLE 1 for ascomycetes. The fast formation of PEVs occurs, but their scission and further endocytosis are blocked. This result indicates the possible involvement of F-actin in the regulation of PMT in basidiomycetes, but the mechanism for such involvement is unknown.

The aim of this study is to develop a model of PMT regulation in filamentous basidiomycetes based on an analysis of: (i) the API effect on the early stages of endocytosis; (ii) the impact of hyperosmotic shock on early endocytosis; (iii) the effects of API and hyperosmosis on plasmolysis and the hyphae sizes. In the study, strains of *R. solani* and *S. hirsutum* that belonging to different ecological trophic groups were used. Having studied in detail the effects of different API concentrations and hyperosmotic condition on the dynamics and micromorphology of fungal endocytosis labeled with fluorescent styryl probe, and on protoplast and hyphal shrinkage, the curtain model of PMT regulation was offered. The model was built without the use of fluorescence or ultrastructural labeling of F-actin, because modern methods of actin labeling in basidiomycetes do not allow obtaining, especially in non-apical cells and especially for actin cables, the required image quality (Salo et al. 1989; Roberson 1992; Gorfer et al. 2001; Timonen, Peterson 2002; Takeshita, Fischer 2019).

## MATERIALS AND METHODS

### Strains and culture media

The main subject of the present research is heterobasidial phytopathogen *R. solani* J.G. Kühn. Some experiments are also duplicated on xylotrophic basidiomycete *S. hirsutum* (Willd.) Pers. Strains of both fungi were used in our previous studies: *R. solani* in Kamzolkina et al. (2017); *S. hirsutum* in Mazheika et al. (2020).

Strains for cytochemical analysis were cultured on Czapek medium with 0.2% NaNO_3_ and 2% agar. Peripheral mycelium of colonies not older than 4–5 days after seeding was used for research. Liquid (agar-free) Czapek medium was used to incubate mycelium in API and sorbitol.

### Labeling of endocytic compartments, the use of APIs and hyperosmotic conditions

Fluorescence labeling of PEVs and other endocytic structures was carried out using an AM4-64 styryl probe (Biotium, Fremont, California). AM4-64 is a full analog of FM4-64FX, which is provided by Molecular Probes/Thermo Fisher. The method for the preparation of mycelium samples and their labeling is given in Kamzolkina et al. (2017) and Mazheika et al. (2020). The same works describe a method for assessing the rate and dynamics of endocytosis. Briefly, the samples were analyzed under a microscope at two time intervals: from 0 min (start of endocytosis with the transfer of samples from ice to room temperature) to 15 min and from 15 to 30 min. The entire common endocytic pathway was divided into four conditional discrete stages, designated from 0 to 3. The number of mycelial cells corresponding to stages 0 to 3 was counted (about 100–150 cells for each slide). In stage 0, AM4-64 labels only the plasmalemma and no PEVs (FIG. 1A); in stage 1, PEVs appear (FIG. 1B); in stage 2, vesicles larger than PEVs (endosomes, scissored macrovesicles) are visible in the cytoplasm, mainly near the plasmalemma (FIG. 1C); and in stage 3, the label appears in the vacuole tonoplasts, including in the large vacuoles (FIG. 1E). Diagrams were constructed, the columns of which reflect the proportion of cells corresponding to one or another of the four stages of endocytosis. Distributions were compared using the chi-square method and were considered statistically different at p < 0.01.

**Figure 1.**
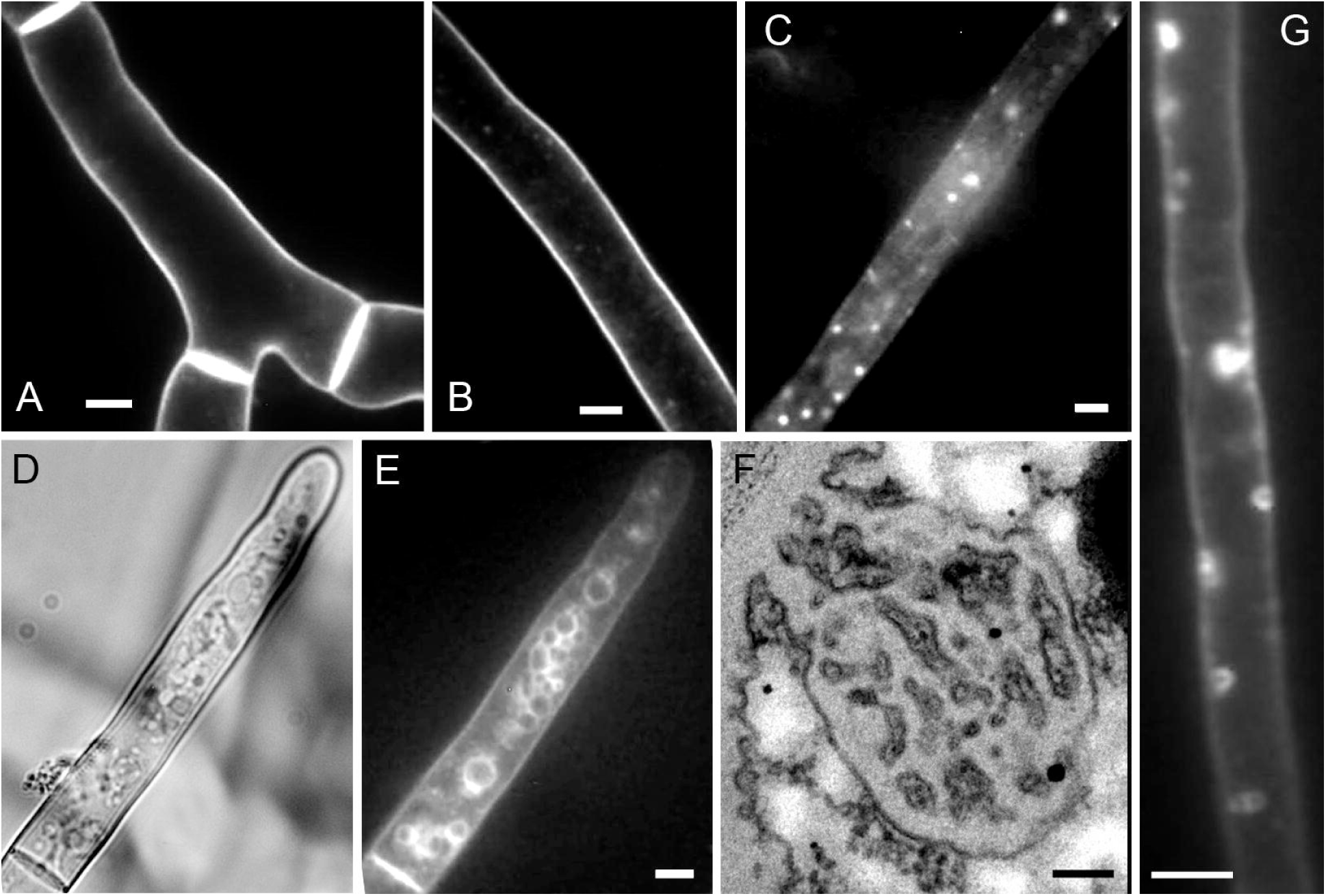
Endocytosis without treatment of mycelium by APIs in *R. solani* (A–E) and *S. hirsutum* (F, G). Microphotographs with fluorescence AM4-64 labeling (except F, showing a TEM photograph of a plicated macrovesicle). A–C, E. Stages of endocytosis from 0 to 3 are sequentially depicted (D is the same site of the slide as E but in transmitted light). G. Stage 1 with macrovesicles in *S. hirsutum*. Bars = everywhere 5 μm, except F, 100 nm.

To inhibit the assembly of actin filaments, mycelium samples were incubated for 20 min at 10, 50, and 100 μM cytochalasin D (CytD; Enzo, Lausen, Switzerland; from 5 mM stock in DMSO: dimethyl sulfoxide; Sigma-Aldrich, St. Louis, Missouri) before using AM4-64. Another variant is incubation in 0.5, 2, 20, 80, and 200 μM latrunculin A (LatA Santa Cruz Biotechnology, Dallas, Texas; from 2.5 mM stock in DMSO).

To study the effect of hyperosmosis on endocytosis, mycelium of *R. solani* was incubated in 0.6 M sorbitol (Sigma-Aldrich). Two treatment options were used: 30 min preincubation in sorbitol before AM4-64 labeling and introduction of mycelium in sorbitol immediately after washing off AM4-64 (at 0 min). In addition, the effect of 0.6 and 1.2 M sorbitol (15 min preincubation before microscopy) with and without APIs on plasmolysis in mycelial cells and on hyphae diameter was documented using transmitted light microscopy. The influence of conditions on plasmolysis was quantified by the ratio of cells with certain types of plasmolysis to the total number of cells (calculated by septa) in the microscopic photos. The septa were visualized using Fluorescent Brightener 28 (FB28, Sigma-Aldrich; final concentration of 40 μg/ml from stock 20 mg/ml in DMSO). The Whitney-Mann test (U-test) was used to statistically evaluate the effect of APIs and hyperosmosis on plasmolysis and hyphae diameter and statistical differentiations were considered at p < 0.01.

Axioskop 40 FL and Imager M epifluorescence microscopes (Zeiss, Jena, Germany) were used in this work.

### Electron microscopy and software

The sample preparation for transmission electron microscopy was conducted according to the previously described protocols (Matrosova et al. 2009).

axiovision le 4.8 (Zeiss) and photoshop 8 (Adobe Systems, San Jose, California) were used to obtain and process microscopic images. For statistical calculations, the software package statistica 6 (StatSoft, Tulsa, Oklahoma) was used.

## RESULTS

### Differences between endocytosis in R. solani and S. hirsutum and effect of CytD on endocytosis in R. solani

FIG. 1 demonstrates the early stages of normal (without treatment by APIs) endocytosis in *R. solani* (FIG. 1A, B) and in *S. hirsutum* (FIG. 1F, G). Endocytosis in *R. solani* and in *S. hirsutum*, as previously shown (Kamzolkina et al. 2017; Mazheika et al. 2020), is active in many cells (not only apical) of the peripheral part of a young colony cultivated on an agarized nutrient medium. FIG. 1 shows that the most important distinction between early endocytosis in *S. hirsutum* and *R. solani* is the absence of macrovesicles in the latter; PEVs in *R. solani* have a smaller scatter in size than in *S. hirsutum* and the largest PEVs are visualized as small points of the fluorescent signal.

The results of a semi-quantitative analysis of the effects of 10, 50, and 100 μM CytD on endocytosis of *R. solani* are shown in the diagrams in FIG. 2A–F. The dynamics of endocytosis are not affected by 10 μM CytD in the observation interval of 15–30 min (FIG. 2A). In the interval 0–15 min, a certain acceleration of PEV scission from the plasmalemma occurs: in FIG. 2A (left-hand diagram), it can be seen that the stage 2 column in the experiment diagram is higher than in the control, and the stage 1 column, by contrast, is lower.

**Figure 2.**
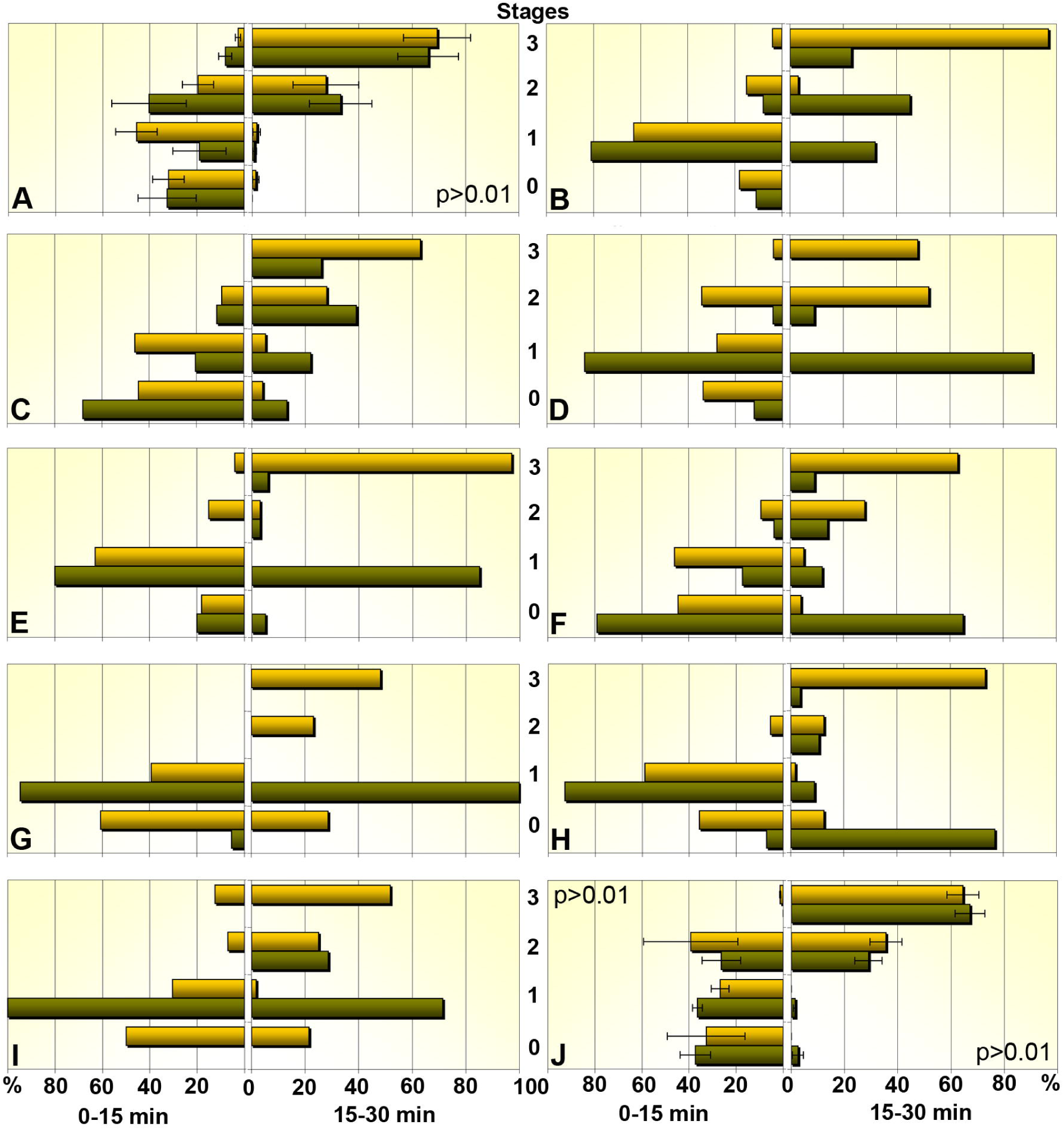
The dynamic of endocytosis of *R. solani* with different treatments is illustrated in the diagrams. Each diagram comprises two parts: the left section corresponds to the observation time interval of 0–15 min, the right to 15–30 min. The stages of endocytosis from 0 to 3 are plotted on the Y-axis; on the X-axis is the occurrence frequency of cells with the corresponding stage of endocytosis in %. Dark columns are according to treated samples, light, to controls. A. Mycelium was treated with CytD at a concentration of 10 μM; averaged data are presented. B–D. Three separate experiments with the processing of mycelium by CytD 50 μM. E, F. Two separate experiments with CytD 100 μM. G–I. Three separate experiments with LatA 2 μM. J. Effect on endocytosis of 0.6 M sorbitol, 30 min pretreatment before labeling by AM4-64, averaged data. For the distributions that are not statistically different in chi-square, p > 0.01 is indicated; in the diagrams without indicated p, p << 0.01.

On *R. solani* endocytosis, 50 μM CytD has a significantly greater and somewhat opposite influence than its 10 μM concentration. In the interval 0–15 min, CytD inhibits the early stages of endocytosis. There are individual diagrams for three independent experiments in FIG. 2B–D (here and below there are not averaged, but diagrams of individual experiments – this approach allows to better understand the biological effect of APIs on endocytosis). In two of these experiments, suppression of PEV scission exists: the frequency of occurrence of cells with stage 1 reaches 80–85% and 5–10% with stage 2 (in the control 15–35%; FIG. 2B, D). In the third experiment (FIG. 2C), PEV formation is slowed down: almost 70% of the cells correspond to stage 0 and 20% to stage 1 versus about 45 and 50% in the control, respectively. In the observation interval of 15–30 min, 50 μM CytD in one of the experiments significantly inhibited endocytosis at stage 1 (about 90% of the cells correspond to stage 1; in the control at this stage, all PEVs were already scissored; FIG. 2D). In two other experiments, endocytosis continued but clearly lagged behind the control (FIG. 2B, C).

The dynamics of *R. solani* endocytosis are similarly affected by 100 μM CytD as by the 50 μM concentration but in a more pronounced manner. In the 0–15 min interval, endocytosis inhibition also occurs at the stages of either the formation or scission of PEVs (FIG. 2E, F, left-hand diagrams). In the late phases of observation, the strong inhibitory effect of CytD persists, unlike the situation with 50 μM concentration, in which, in some experiments, endocytosis, although slowed, reaches stages 2 and 3; this does not happen with 100 μM (FIG. 2E, F, right-hand diagrams).

A qualitative fluorescence microscopic analysis of the effects of 50 and 100 μM CytD on endocytosis of *R. solani* is shown in FIG. 3A–H. The results of morphological analysis are as follows: (i) CytD can, indeed, inhibit PEV scission: the plasmalemma of many mycelial cells, including in the observation interval of 15–30 min (FIG. 3G, H), is covered with vesicles, and the AM4-64 labeled membranes are rare in the cytoplasm. (ii) Attached to the plasmalemma, PEVs increase in size relative to control stage 1; occasionally, small “macrovesicles” can be found (FIG. 3F, arrowhead), not characteristic of the native *R. solani*. (iii)CytD stimulates the formation of tubular invaginations up to 4–5 μm long (FIG. 3C–F, white arrows); PETs are more likely to occur in the observation interval of 0–15 min and more often at 50 μM CytD concentrations than at 100 μM. It is sometimes possible to observe PETs with “heads” (FIG. 3C). (iv) In the late phases of observation, especially when using 100 μM CytD, cells with clearly degrading parietal PEVs are found (FIG. 3H).

**Figure 3.**
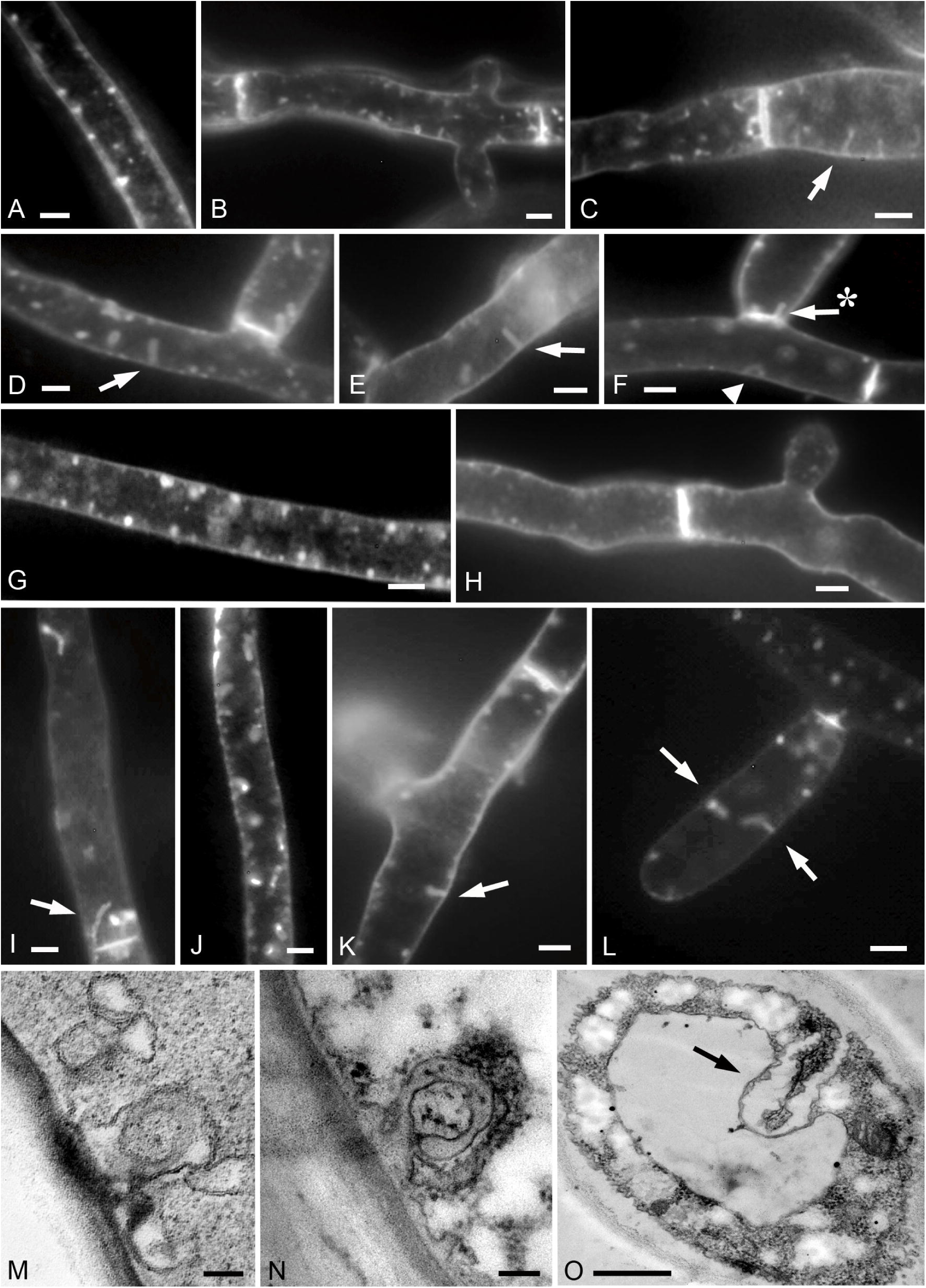
The morphological effect of CytD and LatA on endocytosis of *R. solani* (A–N) and *hirsutum* (O). A–L. Fluorescence microscopic photographs of mycelial cells, endocytotic compartments of which are labeled by AM4-64. M–O. TEM photographs of ultrathin sections through mycelial cells. A–C. Mycelium treated with 50 μM CytD, observation interval 0–15 min; the arrow indicates PET with a “head” (most likely bending of the tubular invagination in the focal plane). D–F. 100 μM CytD, 0–15 min; arrows show tubular invaginations 4–5 μm long; arrow with asterisk shows PET on the septum; the arrowhead labels a large macrovesicle stimulated in *R. solani* by CytD (but it may be bending of the tubular invagination too). G, H. Observation interval 15–30 min; G: 50 μM CytD; H: 100 μM CytD. I–L. Mycelium pretreatment by LatA, observation interval 0–15 min. LatA concentrations from left to right, respectively, are 0.5 μM, 2 μM, 20 μM and 80 μM. Arrows show tubular invaginations. M. U-shaped invagination (with a wide neck, or PET merged with its own base) by the action of 100 μM CytD. N, O. Curved tubular invaginations under the influence of 50 μM CytD in *R. solani* (N) and under the influence of 10 μM CytD in *S. hirsutum* (O, black arrow). It is possible that the rarer occurrence of PETs in *S. hirsutum* compared to *R. solani* is due to the bending of the tubes; in fluorescent preparations, such folded PETs will be indistinguishable from the macrovesicle. Bars = on fluorescence photos 5 μm; M, N: 100 nm; O: 500 nm.

FIG. 3M and 4N show the impact of 100 and 50 μM CytD, respectively, on the ultrastructure of the PEVs in *R. solani*. In TEM samples, PETs bent on themselves are best preserved.

### The effect of LatA and sorbitol on the endocytosis of R. solani and the influence of APIs on endocytosis of S. hirsutum

FIG. 2G–I demonstrates semi-quantitative analysis diagrams showing the effect of 2 μM LatA on the endocytosis of *R. solani*. It can be seen from the diagrams that the inhibitory effect of LatA is similar to CytD, but this inhibitor acts even at a concentration of 2 μM stronger than CytD at 100 μM. In the 0–15 min observation interval, stage 1 almost completely dominates (90–100%), and PEV formation occurs faster than in the control (this is also true for CytD). At the late phase of observation in the two experiments, the vast majority of PEVs remains unscissored (FIG. 2G, I, right-hand diagrams). In the third experiment, there is a return to stage 0 (FIG. 2H, right-hand diagram), which never occurs when CytD is used.

Morphological analysis of fluorescence-microscopic photos of *R. solani* mycelia cells treated with 0.5, 2, 20, and 80 μM LatA (FIG. 3I–L) demonstrates that there are no principled distinctions in the effect on endocytosis of different LatA concentrations. The inhibitory action of LatA also morphologically manifests itself similarly to the action of 50 and 100 μM CytD: there are also many cells with unscissored PEVs; they also increase in size, and the formation of tubular invaginations is also stimulated (in general, the frequency of PET occurrence is somewhat lower than using CytD, but there are invaginations longer than 5 μm; see FIG. 4I, arrow). We did not find significant morphological differences between the effects of 0.5–80 μM LatA concentrations and 200 μM LatA (data not shown).

**Figure 4.**
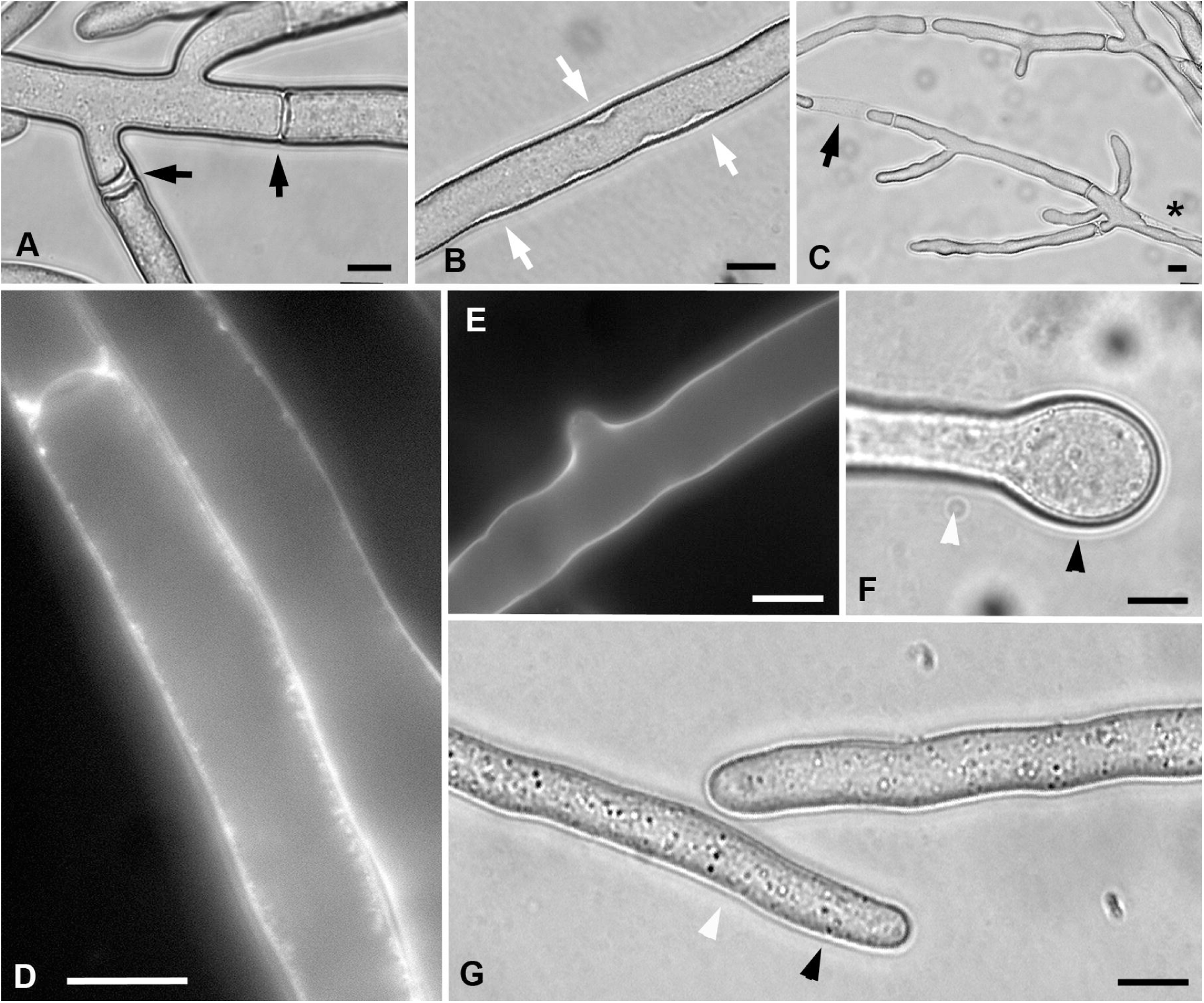
Plasmolysis under hypertonic conditions in *R. solani* cells and the effect of API on hyphal apexes. All photos except D and E are obtained in transmitted light. A, B. Incubation of mycelium in 0.6 M sorbitol; black arrows indicate areas near septa with weak convex-type plasmolysis; white arrows – concave-type plasmolysis sites. C. Exposure to 1.2 M sorbitol; arrow – strong convex-type protoplast shrinkage; asterisk – shrinkage of the protoplast with strong detachment from the side cell walls. D, E. Fluorescent FB28 labeling of cell walls and septa; unlike control (E), the hyperosmotic treatment makes the inner surface of the cell wall rough (D). F, G. Hyphae subapex swelling with 20 μM LatA (F), and control (G); black and white arrowheads mark the diameter measurement sections to quantify the degree of hyphae tip swelling. Bars = 10 μm.

The influence of 0.6 M sorbitol on the dynamics of endocytosis of *R. solani* is reflected in the diagrams in FIG. 2J (diagrams are given for the variant with 30 min preincubation in sorbitol). We did not find an impact of sorbitol on the rate of endocytosis either in the experiments with preincubation in sorbitol or in the experiments with the placement of mycelium in sorbitol after washing off the AM4-64.

In the case of *S. hirsutum*, it is showed that, even when using CytD at a concentration of 10 μM, the scission of PEVs, including macrovesicles, from the plasmalemma is strongly inhibited (data not shown). This result indicates that the sensitivity of *S. hirsutum* endocytosis to actin depolymerization is higher than that of *R. solani*. In general, the number of formed PEVs and, in particular, PETs in *S. hirsutum* is lower than that in *R. solani* treated with 50 or 100 μM CytD. FIG. 3O (arrow) shows a TEM photo of a tubular invagination bent on itself in *S. hirsutum*.

### The effect of API and hyperphase on plasmolysis and mycelial size of R. solani

Table 1 demonstrates that only a small fraction of the cells (about 16%) of *R. solani* mycelium is prone to plasmolysis in 0.6 M sorbitol. In most cases, plasmolysis in these cells is weak, convex-type (Oparka 1994) – the protoplast only slightly shrinks near the septum (FIG. 4A). Concave-type plasmolysis (Oparka 1994) occurs in about a quarter of plasmolytic cells, with the formation of small pockets of plasmalemma along the side walls of hyphae (FIG. 4B). Incubation of mycelium in 1.2 M sorbitol leads to a significant increase in plasmolysis. In many cells, the protoplast does not only strongly shrink at the ends of the cell, but also separates from the side walls of the hyphae (FIG. 4C), which does not occur in 0.6 M sorbitol. The combined use of sorbitol and 20 μM LatA is insignificant, but still affects the plasmolysis of *R. solani*, when compared with the action of only sorbitol (see Table 1). In hyperphase, in many cells of the FB28-labeled mycelium, the inner surface of the cell wall becomes roughened and has FB28 point signals similar to the actin patches (FIG. 4D, E).

**Table 1.**
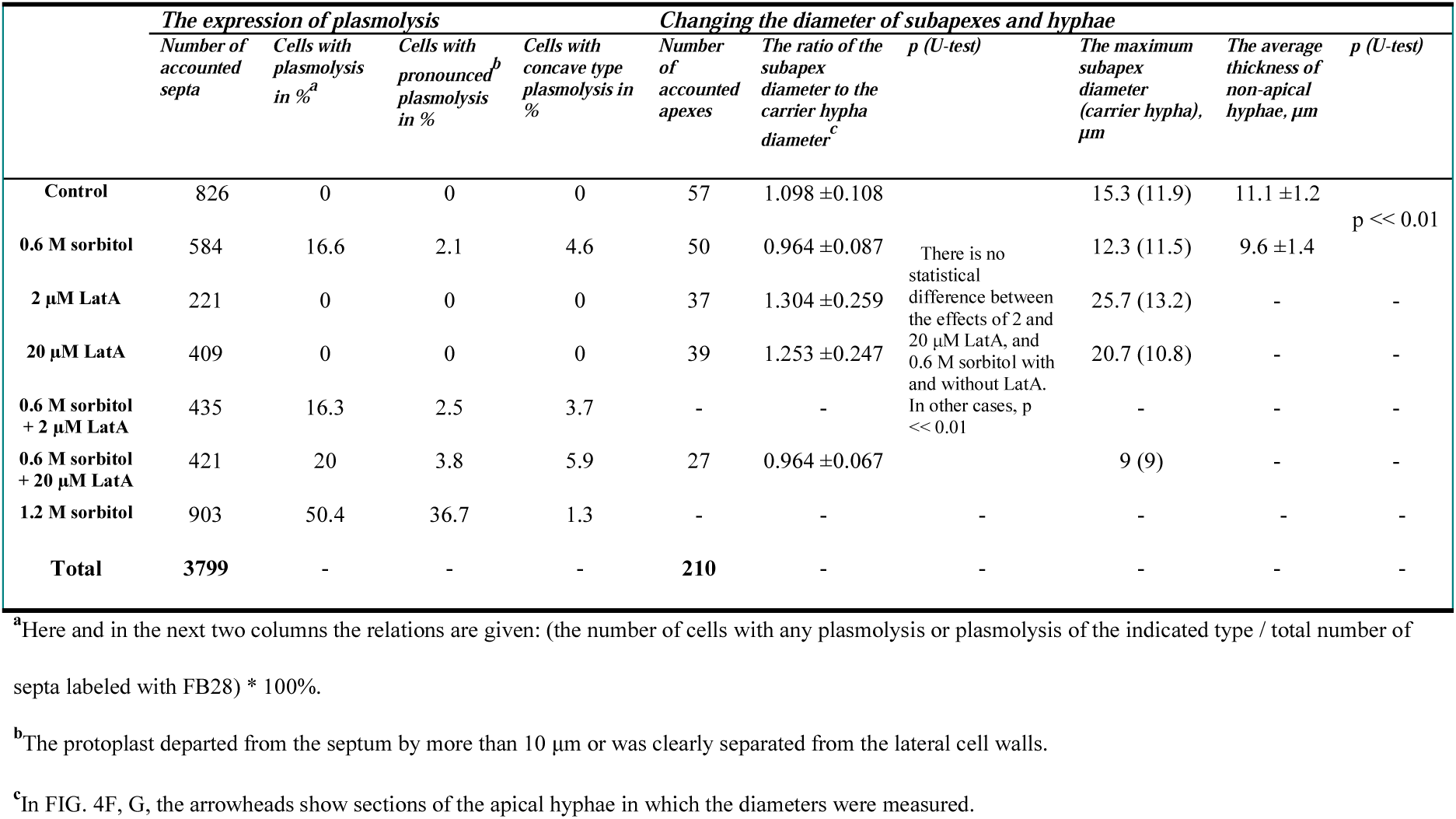
The expression of plasmolysis under different conditions in *R. solani*, the influence of hyperosmosis and LatA on the swelling of apexes/subapexes and on the thickness of the hyphae

It was found that incubation of mycelium in sorbitol leads to a decrease in the diameter of hyphae (Table 1). On average, the thickness of non-apical hyphae in 0.6 M sorbitol decreases by about 15% relative to the control. This difference is statistically significant and it became possible to establish it due to the relative uniformity of *R. solani* hyphae in size (what *S. hirsutum* doesn’t have). Another result associated with resizing hypha relates to hyphal apexes (see Table 1 and FIG. 4F, G). The use of even 2 μM LatA leads to swelling of the subapical zone of many hyphae. In some cases, the tip of the hyphae increased almost two times in diameter relative to the carrying hyphae. An increase in LatA concentration to 20 μM does not give an enhancement of the effect. In the control, some apexes are also slightly swollen (but not in sorbitol), apparently due to the slight hypoosmoticity of the liquid medium to the conditions of cultivation on a solid substrate.

## DISCUSSION

*R. solani* has a patch type of endocytosis like in *Aspergillus*, yeasts, etc. Accordingly, the type of endocytosis inherent in a fungus more depends not on its systematic position, but on belonging to the certain ecological trophic group. *S. hirsutum* has macrovesicular endocytosis that may serve as an internal digestive system, capturing large volumes of extracellular fluid with nutrients by macrovesicles. Thus, xylotrophic and some other basidiomycetes control the microorganisms of their own hyphosphere, preventing them from excessive intercepting hydrolysis products that are formed during exoenzymatic utilization of the substrate by basidiomycete. *R. solani*, being a phytopathogenic fungus, does not experience serious competition for resources with microorganisms, but for it, strictly controlled interaction with the host plant is important. Apparently, the patch type of endocytosis is better suited for this kind of biological interaction.

The fact that the formation of PEVs (and especially PETs) occurs at any LatA concentration (even at 200 μM, at this concentration F-actin is completely depolymerized according to the literature; Basu et al. 2014) indicates that nonapical hyphae have not high turgor pressure (as in many filamentous fungi, SUPPLEMENTARY TABLE 1) and turgor weakly affects PMT. The absence of the influence of 0.6 M sorbitol on the dynamics of early endocytosis in *R. solani* can also serve as confirmation of weak PMT dependence from turgor. Judging by the results, the certain degree of F-actin depolymerization leads to a decrease in PMT, and plasmalemma invaginates due to the action of BAR and other proteins without the need for F-actin, as described in the Introduction.. However, intact F-actin is required to PEV scission from plasmalemma, therefore the certain degree of actin depolymerization leads to blocking the scission of the formed PEVs and PETs. Thus, in many filamentous basidiomycete cells, PMT is determined by the actin cytoskeleton rather than turgor pressure. On the other hand, in single cases, 50 and 100 μM CytD (the destruction of F-actin at these concentrations, of course, is more pronounced than at 10 μM CytD, but weaker than at 2 μM LatA) inhibits the formation of PEVs, not only scission. This result can be explained by the fact that in some experiments turgor pressure in the mycelium was on average slightly higher and also participated in the maintenance of PMT but still as an additional factor to the cytoskeleton, judging by the result that 2 μM LatA never inhibits the formation of PEVs.

One of the explanations why the destruction of F-actin leads to a decrease in PMT and to the rapid formation of plasmalemma invaginations could be permeabilization of the plasmalemma under the action of API. In this case, PMT could be controlled by turgor, but damage to the membrane would lead to a decrease in turgor pressure and, accordingly, in the PMT level. However, it was previously shown that in experiments with propidium iodide, at least 2 μM LatA does not damage the plasmalemma (Kamzolkina et al. 2017), and in this study a violation of the plasmalemma would lead to rapid accumulation of AM4-64 in vacuolar tonoplasts already in the 0-15 min interval, which does not happen. It is possible that the permeability of the plasmolemma does not change to large molecules, but to low molecular weight compounds (ions, for example). Then turgor may fall without a positive reaction of the permeabilization markers. However, the absence of the influence of 0.6 M sorbitol on the dynamics of endocytosis, as before, contradicts the dominance of turgor in the regulation of PMT.

The different sensitivity of early endocytosis in *R. solani* and *S. hirsutum* to API can be explained by the presence of different types of endocytosis in them. In *R. solani*, apparently, all or part of PEVs are dressed with actin scaffold, as are in the patches in ascomycetes. The actin patch delays the formation and scission of PEV to some extent, so its slight softening with 10 μM CytD accelerates early endocytosis. Macrovesicles either do not have the actin coating, or it is not as rigid and powerful as in the actin patches. In addition, the actin cytoskeleton of *S. hirsutum* may for some reason lose its function of PMT regulating at lower API concentrations, therefore, 10 μM CytD does not accelerate early endocytosis, as in *R. solani*, but already blocks PEV scission.

Like in other fungi (see SUPPLEMENTARY TABLE 1), mycelium of *R. solani* has also cells with relatively high turgor pressure. An example is the apical hyphal cells, which swell under the pressure during the destruction of F-actin. However, although probably PMT here is defined more by turgor, the role of the actin cytoskeleton, which retains the shape of the hyphal apex, is also important, since the young thin cell wall cannot withstand turgor pressure without the not damaged actin cytoskeleton.

Certainly, in *R. solani*, the plasmalemma adheres to the inner layer of the cell wall, apparently through FACs. The number of FACs is supposedly higher on the lateral sides of cells than in regions of septa (FIG. 5A, B), judging by the fact that protoplast during plasmolysis primarily departs from septa. The adhesion and the actin cytoskeleton seem to perform to some extent separate functions: F-actin controls PMT, and FACs retain the protoplast shape even at low PMT (FIG. 5C). The weak effect of APIs on plasmolysis (which means that actin is not the main support for maintaining protoplast form) and the rapid response of endocytosis to APIs (which means that not FACs but actin affects PMT) favor the fact that the functions are distributed that way. However, presumably the adhesion and F-actin constitute a single coherent system (see a curtain model below). An additional mechanism has been discovered that mitigates the effects of plasmolysis and prevents protoplast damage – shrinkage in the cross section of the mycelial cells as a whole along with the cell wall (Table 1; FIG. 5D). A similar mechanism is known in yeast (SUPPLEMENTARY TABLE 1).

**Figure 5.**
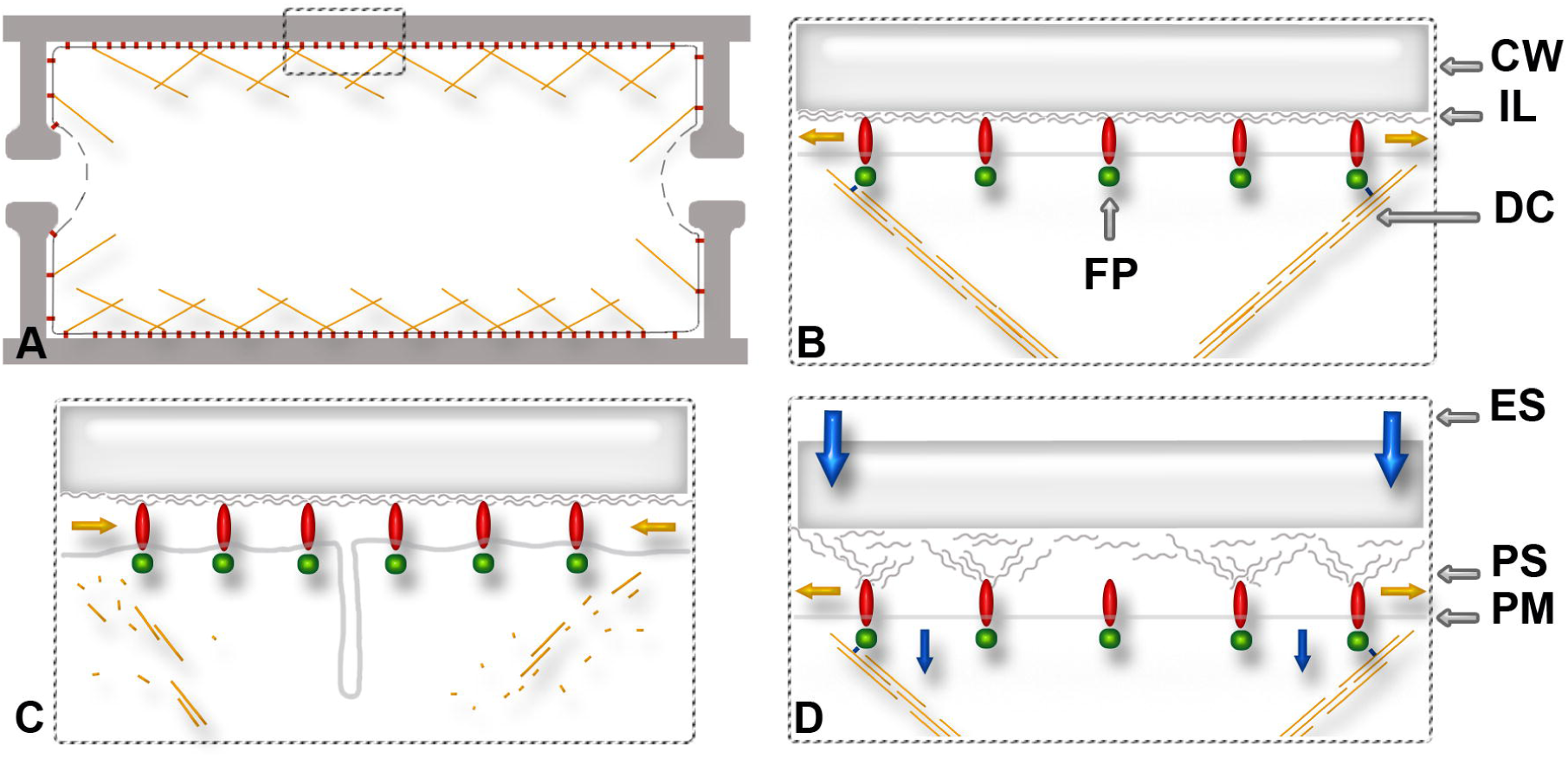
The curtain model of PMT regulation in nonapical hyphal cells in filamentous basidiomycetes. A. Schematic representation of a filamentous basidiomycete cell with the system of actin driver cables regulating PMT. B. Enlarged fragment marked with a dotted frame on A. CW is the cell wall. IL is the inner layer of the cell wall to which FAC proteins are attached. DCs – the actin driver cables, they are connected (via myosin?) with protein complexes (ball in FP) interacting with the intracellular domains of FAC proteins. DCs are associated only with some FACs, due to the motor function of myosin, changes in the cable length or contractile activity of the cables, increase or relaxation of PMT occurs (the mechanism is similar to pushing curtains in and out using curtain drivers – sticks with which curtain rings are moved along the curtain rod). FP – FAC proteins, oval denotes a transmembrane integrin-like protein (or other protein with the same function) that binds the plasmalemma and intracellular elements to the cell wall. C. Hypothesized effects of the API on the fungal cell: the actin driver cables are damaged and cannot carry out the driver function, PMT (with not high turgor pressure in cell) decreases, BAR and other proteins form PEVs and PETs. D. The effect of hyperosmotic treatment of mycelium (0.6 M sorbitol, a cell without the pronounced plasmolysis is shown). The whole hyphae is shrinked in diameter (an average of 15%, large vertical arrows). The periplasmic lumen increases due to the shrinkage of the protoplast (small vertical arrows); however, the proteins of many FACs continue to retain the plasmalemma associated with the cell wall. Due to the force pulling the FACs away from the cell wall, the inner layer of the cell wall loosens, driven by FAC proteins. This explains the roughness of the inner side of the cell wall in FIG. 4D, and capture of “the cell wall material” in PEVs described for yeast and plants (SUPPLEMENTARY TABLE 1). It is important that with moderate hyperosmotic shock, the actin driver cables are not depolymerized (for many fungi the opposite is shown, see SUPPLEMENTARY TABLE 1, perhaps only certain cables are disassembled or in certain cells), which allows to maintain PMT and the previous dynamics of endocytosis.

We offer a curtain model that describes the regulation of PMT in nonapical and non specialized cells of filamentous basidiomycetes (see FIG. 5A–D). According to this model, the plasmalemma, like a curtain, is attached to the cell wall (a curtain rod) through movable FACs (rings on the curtain rod). The certain actin cables in this case serve as curtain drivers (FIG. 5A–B). In the curtain model, it is important that the integrin-like or other proteins in the FACs with their extracellular domains are not rigid anchored to the inner layer of the cell wall, but are able to glide to some extent along the inner surface of the cell wall. If FACs have this property, then regulation of PMT is carried out due actin cables are attached to the intracellular domains of FAC proteins (directly or via intermediate protein complexes) and create the necessary tension on the plasmalemma, just as moving the curtain drivers can close curtains or push them apart.

The main properties of the curtain model are as follows (a number of additional properties are described in the caption to FIG. 5): (i) In most nonapical hyphal cells in filamentous basidiomycetes, the curtain system for regulating PMT is the main one. In certain cells (for example, in apical cells, haustoria, in germinated spores, etc.), the turgor system of regulation of PMT may dominate. These two PMT regulating systems may complement each other: with significant turgor, the curtain regulation is inactive; with reduced osmotic pressure, the curtain system is quickly activated and takes on the function of applying tension to the plasmalemma. (ii) Neighboring hyphal cells with the same osmotic potential (not high) may have the distinct PMT if necessary (for example, for differentially regulating the rate of endocytosis in neighboring cells) which is not available if there is only the osmotic regulatory system that can produce only gradient pressure changes. (iii) The curtain system can provide local PMT reduction in a certain area of the cell surface (for example, at the locus of the formation of a large macrovesicle) without changing the whole cell PMT. This option is also not available with the turgor control system. (iv) Perhaps the actin driver cables are contractile; they move the FAC not only through assembly-disassembly of actin elements or myosin-dependent transport via cables, but also because of the rapid contractions in actomyosin, as in animals. This property is consistent with the work of Reynaga-Peña and Bartnicki-Garcia (2005), in which it was shown that the cytoplasm of apical cells of a number of filamentous fungi can contract; and it was proposed in Heath and Steinberg (1999) also. (v) The curtain model, to some extent, is a variant of the popular biomechanical model of biotensegrity (Reynaga-Peña and Bartnicki-Garcia 2005).

This work, mainly methodologically based on the study of fungal endocytosis, went beyond the limits of the problem of endocytosis only. Through action on turgor and on the actin skeleton, using early endocytosis and plasmolysis as markers, it was shown that PMT, the shape of protoplast, its fit to the cell wall in many nonapical fungal cells depend more on the interaction of the actin cytoskeleton elements with FACs than on turgor pressure. This conclusion is in good agreement with the high ductility of filamentous fungi – the dominance of turgor regulation is excellent for yeast cells or certain mycelial cells but not for most hyphal cells. Such dominance in hyphae would lead to mycelium inertness: hyphae would resemble inflated thin tubes, their physical plasticity would decrease, as well as the rate of physiological adaptation to changing conditions, neighboring cells will not be able to independently regulate many of their processes, etc. Probably, the curtain model can be extended to many filamentous fungi and fungi-like organisms, not only to basidiomycetes.

Unfortunately, the fluorescence methods for labeling F-actin in nonapical cells of basidiomycetes either do not work, or do not provide sufficient image quality (phalloidins, immune labeling). Most likely, the development of methods for labeling actin of filamentous basidiomycetes through genetic transformation (LifeAct and other labeling systems) will also be insufficient for the tasks described. In turn, ultrastructural methods, such as immunogolding, do not allow preserving actin structures to the right extent. Therefore, for the further development of the curtain model of mycelial fungal cell organization, it is necessary to develop effective methods for conserve and labeling fungal F-actin, preferably at the ultrastructural level; and studies are also required with violation of specific protein-protein interactions (e.g., actin bonds with FACs proteins).

## Supporting information

Legend for Supplementary Figure

## ACKNOWLEDGMENTS

The authors would like to thank prof. Oxana L. Kolomiets and the staff of her Cytogenetics Laboratory of Vavilov Institute of General Genetics for their comprehensive support.

## FUNDING

The reported study was funded by Russian Foundation for Basic Research according to the research project № 16-04-00814 № and conducted within the framework of the State Assignment: contracts № 0112-2019-0002 (VIGG RAS) and № AAAA-A16-116021660088-9 (MSU, part 2, section 01 10336). The Moscow University Development Program (UDP-10) provided part of the equipment for this study.

## Notes

### Competing Interest Statement

The authors have declared no competing interest.

### Summary of Updates

Fixed some bugs and inaccuracies

