## Supplementary figures and images for "Early endocytosis as a key to understanding mechanisms of plasmalemma tension regulation in filamentous fungi"

### Supplementary Figure

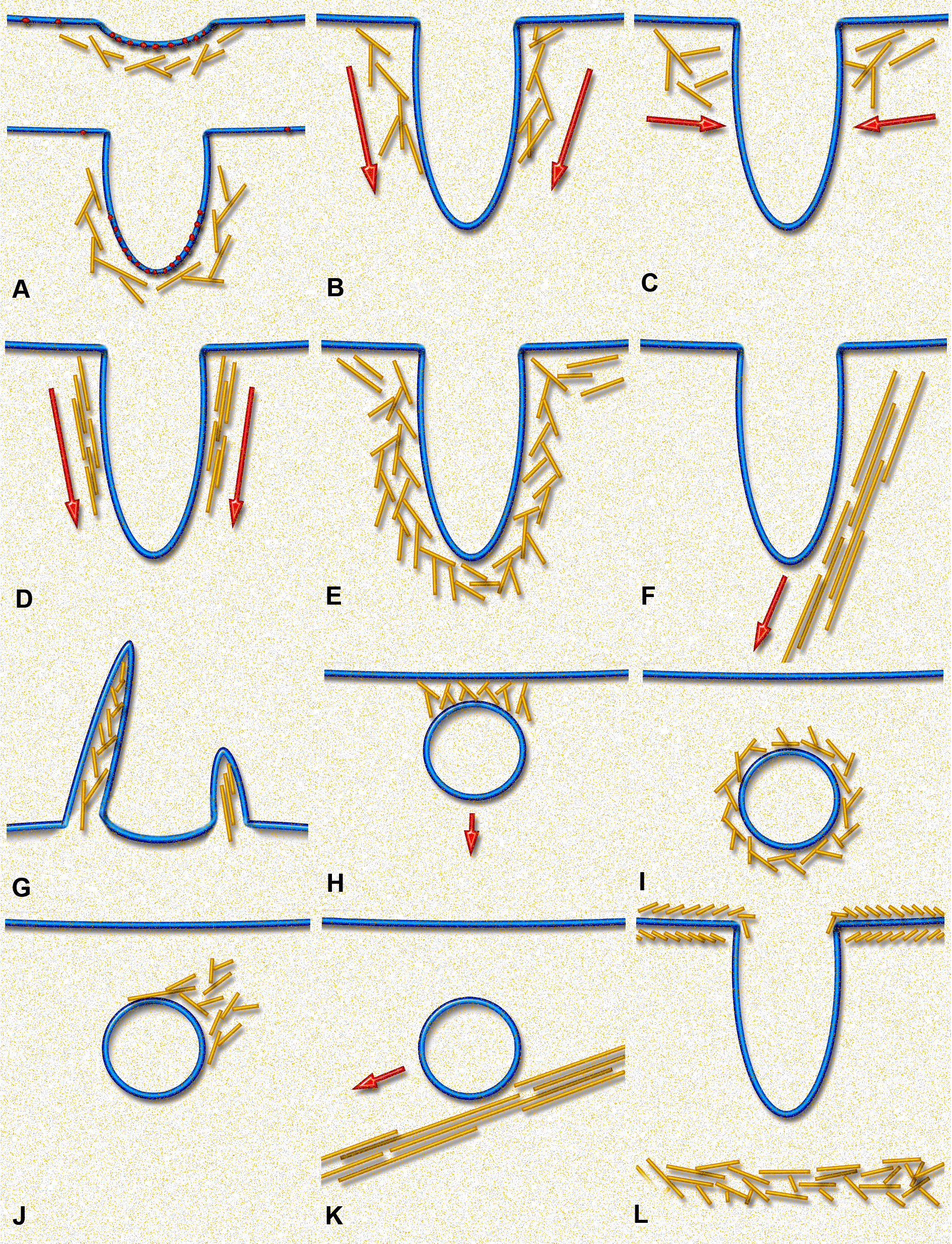
