## Supplementary material for "Early endocytosis as a key to understanding mechanisms of plasmalemma tension regulation in filamentous fungi": Legend for Supplementary Figure

**Supplementary Figure 1.** Possible involvement of F-actin in the early stages of endocytosis in various organisms. The yellow linear elements depict F-actin filaments; their branching is an Arp2/3-actin network; their parallel connection are cables and minicables (usually formin-dependent); both structures are cross-linked to varying degrees (not shown). A–G. Before PEV scission from the plasmalemma. H–K. Immediately following PEV scission. L. No direct involvement of F-actin in endocytosis (the actin cortex is shown at the bottom of panel; its depolymerization and perforation affect the rate of early endocytosis (Itoh et al. 2005; Malacombe et al. 2006; Hinze and Boucrot 2018); at the top: hypothetical peripheral actin networks, intracellular and external). A. Hypothetical mechanism in which actin itself actively clusters specific membrane lipids, provides initial invagination, and then the formation of PEV (Renard et al. 2018). B. The actin network acts on the principle of a strut to elongate the PEV neck. C. The actin network tightens the neck of the PEV. D. Hypothetical analogue of variant B, but the actin minicables, not only the Arp2/3 network, operate (Smythe and Ayscough 2006). E. Fungal actin patch; a strongly cross-linked actin network forms a scaffold around the PEV (Berepiki et al. 2011). F. Actin rails, a hypothetical structure that may be involved in the elongation of the PEV neck and its scission. Rails are described only in animal exocytosis (Malacombe et al. 2006); in endocytosis in FDS, the role of rails is usually attributed to microtubules (Hinze and Boucrot 2018; Renard et al. 2018). G. Classic version of the macropinosome formation in animal cells through protrusion of the plasmalemma to the outside of the cell (Bloomfield and Kay 2016; Marques et al. 2017; Hinze and Boucrot 2018). H. Actin springboard; the myosin-actin network from the bottom of the PEV repels it from the plasmalemma (Hinze and Boucrot 2018). I, J. Two variants for accompanying PEV with actin after scission: scaffold around the vesicle and “comet tail” (there is an opinion that even endosomes inherit the actin scaffold of the PEV) (Huckaba et al. 2004; Malacombe et al. 2006; Moseley and Goode 2006; Berepiki et al. 2011; Mooren et al. 2012). K. Actin cables picking up PEVs after their scission and transporting them to the cytoplasm (Berepiki et al. 2011). L. See above.

**LITERATURE CITED** (only those links that are not in the main text of the paper are given)

Bloomfield G, Kay RR. 2016. Uses and abuses of macropinocytosis. Journal of Cell Science 129:2697–2705

Huckaba TM, Gay AC, Pantalena LF, Yang H-C, Pon LA. 2004. Live cell imaging of the assembly, disassembly, and actin cable-dependent movement of endosomes and actin patches in the budding yeast, *Saccharomyces cerevisiae*. Journal of Cell Biology 167:519–530.

Malacombe M, Bader M-F, Gasman S. 2006. Exocytosis in neuroendocrine cells: New tasks for actin. Biochimica et Biophysica Acta – Molecular Cell Research 1763:1175–1183.

Marques PE, Grinstein S, Freeman SA. 2017. SnapShot: Macropinocytosis. Cell 169:766–766.

Moseley JB, Goode BL. 2006. The yeast actin cytoskeleton: from cellular function to biochemical mechanism. Microbiology and Molecular Biology Reviews 70:605–645.

Smythe E, Ayscough KR. 2006. Actin regulation in endocytosis. Journal of Cell Science 119:4589–4598.
