## Supplementary Table for "Early endocytosis as a key to understanding mechanisms of plasmalemma tension regulation in filamentous fungi"

**Supplementary Table 1.** Turgor pressure and the effect of hyperosmotic shock and of API using on endocytosis and other processes in yeast and in vegetative mycelium of fungi and oomycetes**^a^** (according to literature data)

|  | ***Yeasts*^b^** | | ***Filamentous ascomycetes*^d^** | | ***Oomycetes*^e^** | |
| --- | --- | --- | --- | --- | --- | --- |
|  | ***In hyperosmotic conditions*** | ***Using APIs*** | ***In hyperosmotic conditions*** | ***Using APIs*** | ***In hyperosmotic conditions*** | ***Using APIs*** |
| **Turgor pressure** | According to one of the latest estimates, budding yeast have about 0.2 MPa. May be relatively high, up to 1–1.5 MPa in fission yeast for example | | In general, not relatively high, for example, about 0.5 MPa in *Neurospora crassa*. Most likely, turgor is different in different cells of the same mycelium and depends on the conditions. The apical cell is probably often under a higher turgor. This is very important for the growth of the apex, invasion in the substrate, etc. Turgor falls, and then recovers to its original value under hypershock conditions | | Less dependent on turgor than ascomycetes. Oomycetes can grow at zero turgor. If turgor pressure is present, then it is also relatively not high (for example, about 0.2–0.7 MPa in *Achlya bisexualis*). The apical cell may or may not be under pressure. Growth and invasive ability are dependent more on the actin cytoskeleton. After hyperosmotic shock, turgor may be not restored | |
| **Plasmolysis** | The plasmalemma usually does not separate from the cell wall, the whole cell shrinks, the cell wall instant thickens | **-** | Hyphae are often resistent to plasmolysis; in 0.4–0.8 M sorbitol, the plasmalemma retracts from the cell wall only in individual cells, often only in form of separate pockets. The incipient plasmolysis method of turgor measurement gives overestimated results | - | Same as in filamentous ascomycetes | - |
| **Formation of PEV (PET)** | Yes, numerous PEV and PET across the entire surface of the plasmalemma. They may contain the cell wall material, judging by the staining with calcofluor (a similar result is shown for plant cells). But some or even most invaginations may be deepened eisosomes**^c^** | No, a complete formation blocking. The formation can partially recover when placed in hyperosmotic media | Yes, but few data; they may be large up to about 2 μM | Yes (judging by the photo in a number of publications)**^f^** | No data, but patches appear in the hyphal tips | No data |
| **PEV (PET) scission** | Blocked | **-** | Blocked | Blocked | No data | No data |
| **Actin cables disassembly** | Yes | Yes | Yes | Yes | Yes, disassembling the F-actin cap (actin network in the hyphal tips) | Yes, disassembling the F-actin cap |
| **Growth stop** | Yes | Yes | Yes | Yes | No data | Yes |
| **Apex/subapex swelling** | **-** | **-** | No or no data | Yes | Yes**^g^** | Depending on the turgor in the hyphal apexes – if there is high pressure, then the subapex swells, if not – no or weak swelling |

**^a^**In filamentous fungi and oomycetes, in most cases, only apical and subapical hyphae cells, or conidia seedlings, were studied.

**^b^**Main yeast objects: *Saccharomyces cerevisiae* and *Schizosaccharomyces pombe* (Morris et al. 1986; Oparka 1994; Slaninova et al. 2000; Aghamohammadzadeh and Ayscough 2009; Smaczynska-de Rooij et al. 2011; Basu et al. 2014; Palmer et al. 2015; Goldenbogen et al, 2016; Elhasi, Blombers 2019).

**^c^**Eisosomes are a kind of functional analogue of caveolae in animal cells, found in yeast and some algae. In filamentous fungi, characteristic foci of the fluorescent signal are detected when tagging proteins for eisosomes (Vangelatos et al. 2010), but the issue is not well studded. Eisosomes are invaginations of the plasmalemma in the form of elongated grooves, formed due to specific BAR proteins. The main difference from any PEVs is that eisosomes do not have scission and do not participate in endocytosis. In hyperosmotic conditions, they deepen to inside a yeast cell (Moseley 2018; Appadurai et al. 2020).

**^d^**Main objects among filamentous ascomycetes: *N. crassa* and *Aspergillus nidulans*, less commonly other ascomycetes species (Amir et al. 1995; Torralba et al. 1998; Heath, Steinberg 1999; Lew et al. 2004; Peñalva 2005; Lew 2010, 2019; Bitsikas et al. 2011; Walker, White 2017).

**^e^**Some model species from Saprolegniaceae and Peronosporaseae (Gupta, Heath 1997; Heath, Steinberg 1999; Chitcholtan, Garrill 2005; Walker et al. 2006).

**^f^**In a number of works using APIs, the authors state the blocking of endocytosis. However, from the published photos it can be seen that such blocking is accompanied by the formation of plasmalemma-linked PEVs (Yamashita and May 1998; Upadhyay and Shaw 2008; Lewis et al. 2009). A similar result was obtained for the basidial xylotroph *Lentinula edodes* (Lee et al. 2007).

**^g^**Walker et al. 2006.

**LITERATURE CITED** (only those links that are not in the main text of the paper are given)

Amir R, Steudle E, Levanon D, Hadar Y, Chet I. 1995. Turgor changes in *Morchella esculenta* during translocation and sclerotial formation. Experimental Mycology 19:129–136.

Appadurai D, Gay L, Moharir A, Lang MJ, Duncan MC, Schmidt O, Teis D, Vu TN, Silva M, Jorgensen EM, Babst M. 2020. Plasma membrane tension regulates eisosome structure and function. Molecular Biology of the Cell 31:287–303.

Bitsikas V, Karachaliou M, Gournas C, Diallinas G. 2011. Hypertonic conditions trigger transient plasmolysis, growth arrest and blockage of transporter endocytosis in *Aspergillus nidulans* and *Saccharomyces cerevisiae*. Molecular membrane biology 28:54–68.

Goldenbogen B, Giese W, Hemmen M, Uhlendorf J, Herrmann A, Klipp E. 2016. Dynamics of cell wall elasticity pattern shapes the cell during yeast mating morphogenesis. Open biology 6:160136.

Gupta GD, Heath IB. 1997. Actin disruption by latrunculin B causes turgor-related changes in tip growth of *Saprolegnia ferax* hyphae. Fungal Genetics and Biology 21:64–75.

Lee MT, Szeto CYY, Ng TP, Kwan HS. 2007. Endocytosis in the shiitake mushroom *Lentinula edodes* and involvement of GTPase LeRAB7. Eukaryotic Cell 6:2406–2418.

Lew RR. 2010. Turgor and net ion flux responses to activation of the osmotic MAP kinase cascade by fludioxonil in the filamentous fungus *Neurospora crassa*. Fungal Genetics and Biology 47:721–726.

Lewis MW, Robalino IV, Keyhani NO. 2009. Uptake of the fluorescent probe FM4-64 by hyphae and haemolymph-derived in vivo hyphal bodies of the entomopathogenic fungus *Beauveria bassiana*. Microbiology 155:3110–3120.

Morris GJ, Winters L, Coulson GE, Clarke KJ. 1986. Effect of osmotic stress on the ultrastructure and viability of the yeast *Saccharomyces cerevisiae*. Microbiology 132:2023–2034.

Moseley JB. 2018. Eisosomes. Current Biology 28:R376–R378.

Palmer SE, Smaczynska-de Rooi, II, Marklew CJ, Allwood EG, Mishra R, Johnson S, Goldberg MW, Ayscough KR. 2015. A dynamin-actin interaction is required for vesicle scission during endocytosis in yeast. Current Biology 25:868–878.

Peñalva MA. 2005. Tracing the endocytic pathway of *Aspergillus nidulans* with FM4-64. Fungal Genetics and Biology 42:963–975.

Smaczynska-de Rooij II, Allwood EG, Mishra R, Booth WI, Aghamohammadzadeh S, Goldberg MW, Ayscough KR. 2011. Yeast dynamin Vps1 and amphiphysin Rvs167 function together during endocytosis. Traffic 13:317–328.

Slaninová I, Šesták S, Svoboda A, Farkaš V. 2000. Cell wall and cytoskeleton reorganization as the response to hyperosmotic shock in *Saccharomyces cerevisiae*. Archives of microbiology 173:245–252.

Torralba S, Raudaskoski M, Pedregosa AM, Laborda F. 1998. Effect of cytochalasin A on apical growth, actin cytoskeleton organization and enzyme secretion in *Aspergillus nidulans*. Microbiology 144:45–53.

Upadhyay S, Shaw BD. 2008. The role of actin, fimbrin and endocytosis in growth of hyphae in *Aspergillus nidulans*. Molecular Microbiology 68:690–705.

Vangelatos I, Roumelioti K, Gournas C, Suarez T, Scazzocchio C, Sophianopoulou V. 2010. Eisosome organization in the filamentous ascomycete *Aspergillus nidulans*. Eukaryotic cell 9:1441–1454.

Yamashita RA, May GS. 1998. Constitutive activation of endocytosis by mutation of myoA, the myosin I gene of *Aspergillus nidulans*. Journal of Biological Chemistry 273:14644–14648.
